# A male infertility mutation reverts NANOS1 activity from anti-apoptotic to pro-apoptotic by disrupting repression of *GADD45A*, *GADD45B*, *GADD45G* and *RHOB* genes

**DOI:** 10.1101/858654

**Authors:** Damian M. Janecki, Erkut Ilaslan, Maciej J. Smialek, Marcin P. Sajek, Maciej Kotecki, Barbara Ginter-Matuszewska, Anna Spik, Patryk Krainski, Jadwiga Jaruzelska, Kamila Kusz-Zamelczyk

## Abstract

**Background:** While Nanos-mediated maintenance of germ cells in *Drosophila* and mice has been related to regulation of apoptosis, the relevance of these findings to human physiology is uncertain. Previously we have described the p.[(Pro34Thr);(Ser83del)] double *NANOS1* mutation as associated with an absence of germ cells in the testes of infertile patients. The aim of this study was to identify the mechanism underlying infertility phenotype of patients bearing the *NANOS1* mutation.

**Methods:** Constructs encoding a wild-type or mutated NANOS1 protein were used for transfection of TCam-2 cell line, representing male germ cells in culture. Influence of this mutation on cell proliferation was performed using MTS assay while apoptosis and cell cycle were measured by flow cytometry. RNA-Seq analysis including quantitative RT-PCR was conducted for selecting pro-apoptotic genes, repressed by the wild-type NANOS1. Influence of the p.[(Pro34Thr);(Ser83del)] NANOS1 mutation on that repression was investigated by quantitative RT-PCR.

**Results:** We show here that the p.[(Pro34Thr);(Ser83del)] double *NANOS1* mutation causes NANOS1 to functionally switch from being anti-apoptotic to pro-apoptotic in the human male germ cell line TCam-2. This mutation disrupts repression of mRNAs encoding pro-apoptotic GADG45A, GADD45B, GADD45G and RHOB factors, which could contribute to an increase in apoptosis.

**Conclusions:** This report underscores the conservation of Nanos from flies to humans as a repressor of pro-apoptotic mRNAs in germ cells, and provides a basis for understanding NANOS1 functions in human reproductive health.

## Introduction

Nanos morphogen plays crucial roles in multiple aspects of *Drosophila* body patterning [1] and germline development, in both sexes [2]. Nanos contains a highly conserved zinc-finger RNA-binding domain in the C-terminal region and acts as a post-transcriptional repressor of specific mRNAs, such as *CycB* (*Cyclin B*) mRNA, in primordial germ cells (PGCs) during their migration to the primary gonads [3]. In particular, Nanos was shown to repress caspase activators such as *hid* (*head involution defective*) and *skl* (*sickle*), to inhibit apoptosis in PGCs. This inhibition is crucial to the survival of PGCs during migration to the primary gonads [4]. In mice, Nanos2 and Nanos3 also play anti-apoptotic roles in PGCs [5]. While NANOS has also been implicated in human germ cell development [6], functions in humans and their underlying mechanisms, such as regulation of apoptosis that has been identified in other species, are still uncharacterized.

Among the three paralogues, *Nanos1*, *Nanos2* and *Nanos3*, that are expressed in mammals, only *Nanos2* and *Nanos3* have been reported to have a role in mouse germ cell development [7–9]. *Nanos2* is necessary for male, but not female germ cell development in mice [9]. *Nanos3* is expressed in bipotential PGCs and thus is important for fertility of both sexes in mice [9]. Unlike the infertility resulting from the knockout of *Nanos2* and *Nanos3*, *Nanos1* knockout mice are viable and fertile, indicating that *Nanos1* is dispensable for mouse development and fertility [10]. Among the three human paralogues, however, only NANOS1 (the paralogue with the highest conservation within human populations [11]) has been identified to be associated with male infertility [12–14]. Namely, a NM_199461.4(NANOS1_v001):c.[100C>A;240_242del]; NM_199461.4(NANOS1_i001):p.[(Pro34Thr);(Ser83del)] double mutation (in this report called the p.[(Pro34Thr);(Ser83del)]) was identified in two patients of Polish origin, manifesting absence of germ cells in seminiferous tubules [14]. It was absent in 400 fertile males of Polish origin and 100 male individuals from other populations. This mutation encompasses the N-terminal conserved NIM-region (**Fig. 1A**), which is necessary for recruitment of deadenylase complex to deadenylate and degrade repressed mRNA targets [15].

**Figure 1.**
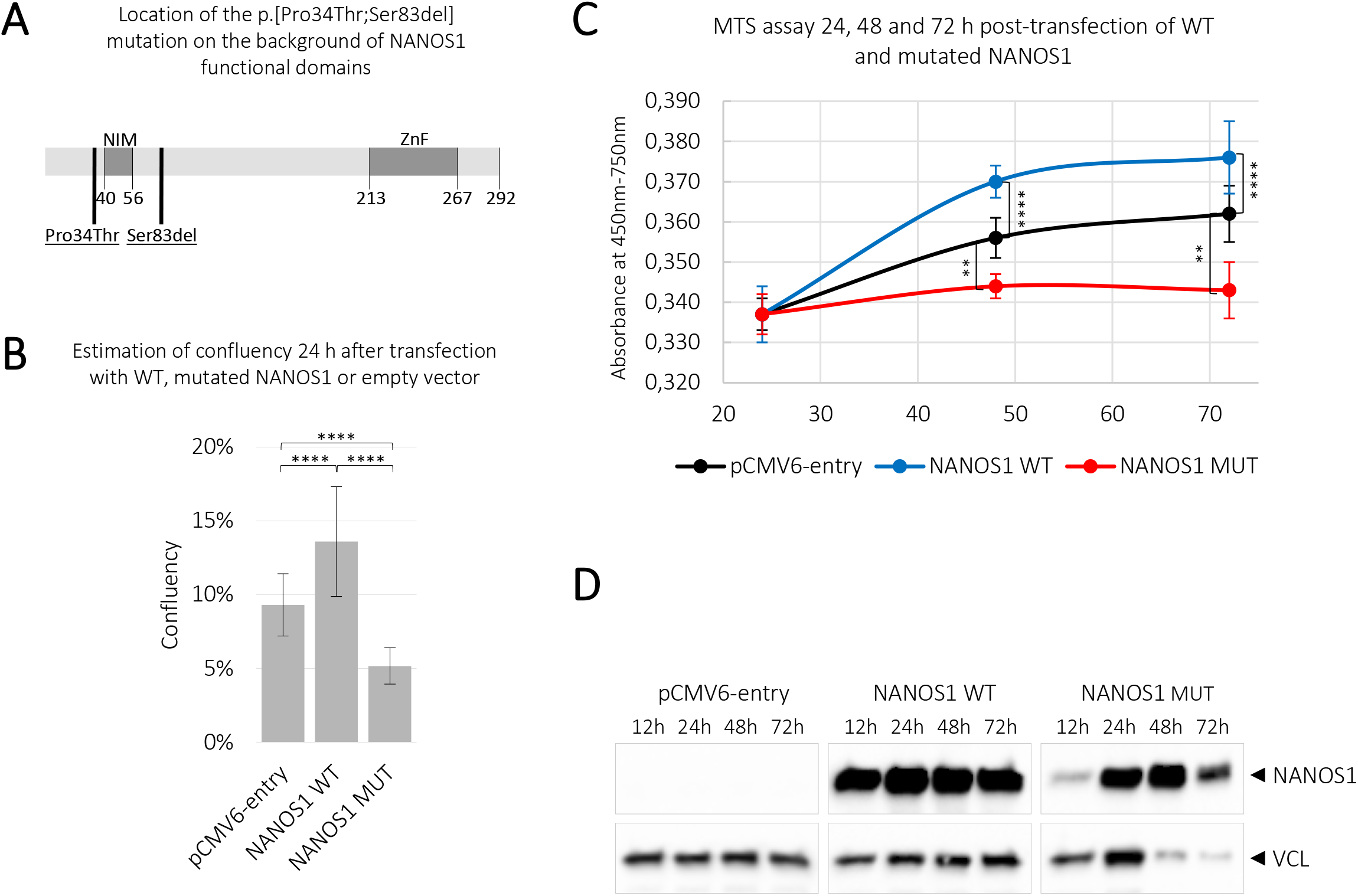
Analysis of the effect of the wild-type and mutated NANOS1 p.[(Pro34Thr);(Ser83del)] on confluency and cell proliferation. TCam-2 cells were transfected with plasmids encoding the wild-type (WT), mutated (MUT) NANOS1 or empty pCMV6-entry vector. **A** – Scheme of the NANOS1 protein. NOT1 interacting motif (NIM) domain, zinc-finger (ZNF) domain and amino acid position of the double mutation p.[(Pro34Thr);(Ser83del)] are indicated. **B** – Confluency was measured 24 h after transfection using JuLI FL, a fluorescence live cell movie analyzer. **C** – The MTS assay was used to analyse cell proliferation of TCam-2 cells 24, 48 and 72 h after transfection. A similar number of cells counted 12 h after transfection was used for normalization of each experiment. This experiment was performed in three biological replicates. **D** – The expression of the wild-type and mutated NANOS1 constructs after transfection compared to the negative control (transfection with empty plasmid) were analysed by western blot analysis using an anti-DDK antibody (OriGene). As loading control VCL (vinculin) protein was used, in each sample shown in lower panel. The *P* value < 0.05 was considered statistically significant, the *P* value < 0.005 was marked by two stars (**) and the *P* value < 0.00005 by four stars (****).

We conducted our study in human male germ cell line, in order to identify the mechanism underlying infertility phenotype of the patients bearing the p.[(Pro34Thr);(Ser83del)] double *NANOS1* mutation. We found that NANOS1 inhibits apoptosis of T-Cam2 cells while repressesing pro-apoptotic genes *GADD45A*, *GADD45B*, *GADD45G* and *RHOB*. Significantly, we also found that this NANOS1 double mutation causes enhancement of apoptosis while disrupting repression of the above pro-apoptotic genes, which might explain the absence of germ cells in seminiferous tubules of infertile patients.

## Materials and Methods

### Cell culture and transfection

TCam-2 cells were obtained from Dr. Sohei Kitazawa and were maintained in RPMI with GlutaMAX medium (Gibco, Life Technologies) supplemented with 10% (v/v) foetal bovine serum (HyClone) and 1% (v/v) antibiotic antimycotic solution (Lonza; Cat. No. 17-745E). The cells were transfected with plasmid constructs using the Neon Transfection System (Life Technologies) according to the manufacturer’s protocol.

### Estimation of cell confluency

2×10^5^ TCam-2 cells were transfected with 1.5 µg of constructs encoding the wild-type, the mutated NANOS1 p.[(Pro34Thr);(Ser83del)] or the empty pCMV6-entry vector (OriGene) in three biological replicates (three independent transfections). Cell confluency was measured 24 h after transfection at 5 random areas of each 6-well plate using JULI FL live cell movie analyzer (NanoEntek).

### MTS proliferation assay

2×10^5^ TCam-2 cells were transfected with 1.5 µg of constructs encoding the wild-type, mutated NANOS1 p.[(Pro34Thr);(Ser83del)] or empty pCMV6-entry vector (OriGene) in three biological replicates (three independent transfections; 5 technical repeats in each on 5 different wells of a 96-well plate) and were cultured for 24, 48, and 72 h after transfection. To obtain similar number per well of viable cells expressing different constructs for MTS assay, viable cells were counted 12 h post-transfection using Trypan Blue staining. Similar number of cells were seeded to 96-well plates. The cell viability was measured using GLOMAX (Promega) at 450 nm (and 750 nm for cell background) wavelength 24, 48, and 72 h after transfection and 4 h after adding 20 µl of CellTiter 96® AQ One Solution Reagent (Promega) to each well containing transfected TCam-2 cells.

The overexpression of the wild-type and mutated NANOS1 was measured 12, 24, 48 and 72 h after transfection by western blot using an anti-DDK antibody (OriGene). VCL (vinculin) protein was used as a loading control.

### Western blotting

Overexpression efficiency was measured by western blot analysis under standard conditions, using a nitrocellulose membrane, horseradish peroxidase (HRP)-conjugated secondary antibodies. Semi-quantitative measurement of protein levels was performed using the ImageLab 5.1 software (BioRad). The chemiluminescent signal was detected using the Clarity™ Western ECL Substrate for HRP (BioRad).

### Antibodies

The primary antibody anti-DDK (FLAG^®^) (OriGene Technologies, TA50011, 1:2500) was used for detection of proteins expressed with the pCMV6-entry vector. Anti-ACTB (actin beta) (Sigma Aldrich, A2066, 1:10000) and anti-VCL (vinculin) (Abcam, ab129002, 1:20000) antibodies were used as loading controls. Secondary antibodies: goat anti-rabbit IgG-HRP (Sigma Aldrich, A6154, 1:250000) and goat anti-mouse IgG-HRP (Santa Cruz Biotechnology, sc-2005, 1:10000) were used.

### Flow cytometry

Detection of apoptotic TCam-2 cells was performed 48 h after transfection of 3×10^5^ TCam-2 cells with 2 µg of constructs encoding the wild-type, mutated NANOS1 p.[(Pro34Thr);(Ser83del)] or empty pCMV6-entry vector. Cells were stained using the Annexin V-FITC Apoptosis Detection Kit (Beckman Coulter) according to the manufacturer’s protocol and measured by flow cytometry using a FlowSight instrument (Amnis). The results were analysed using the Image Data Exploration and Analysis Software version 6.0 (IDEAS^®^ v6.0, Amnis). The experiment was performed in three biological replicates.

Cell cycle analysis was performed 48 h after transfection of 2×10^6^ TCam-2 cells with 20 µg of constructs encoding the wild-type, mutated NANOS1 p.[(Pro34Thr);(Ser83del)] or an empty pCMV6-entry vector co-transfected with GFP-F in proportion 9:1. For this purpose, TCam-2 cells were washed with PBS and fixed in cold 100% methanol on ice for 10 minutes. After incubation at 37 °C for 15 minutes in 50 µg/ml propidium iodide (PI) (Sigma) containing 330 µg/ml RNase A (Sigma), cells were incubated for 1 h on ice and finally measured using the S3e™ Cell Sorter (BioRad) apparatus. Data files were analysed using the ModFit LT (Verity Software House). The experiment was performed in three biological replicates.

### cDNA library preparation and RNA sequencing

2×10^5^ TCam-2 cells were transfected with 1.5 µg of constructs encoding the wild-type NANOS1 or empty pCMV6-entry vector in three biological replicates. Transcription was stopped by Actinomycin D (5 µg/ml) 24 h after transfection for 4 h. Total RNA was extracted using RNeasy Plus Micro Kit (Qiagen) according to the manufacturer’s protocol. RNA quality was checked on Bioanalyzer (Agilent) using RNA 6000 Nano Kit (Agilent). Total RNA samples with RIN value > 7 were used for library preparation. cDNA libraries were prepared using TruSeq RNA Sample Prep v2 (Illumina) and subsequent next-generation sequencing was performed on an Illumina HiSeq 4000 platform by Macrogen INC.

### Bioinformatic analysis

Differential expression analysis was performed on Galaxy platform by the following pipeline: Paired-End sequences obtained from the HiSeq 4000 platform were trimmed using the Trimmomatic v0.36.3 [16]. Sequence reads that passed quality filters were mapped to the human reference genome hg38 using Bowtie 2 v2.3.4.1 [17]. Mapped reads from 3 biological repetitions of NANOS1 and empty vector overexpression libraries were counted using featureCounts v1.6.0.3 [18] followed by calculating differential gene expression with DESeq2 v2.11.40.1 [19]. Differentially expressed transcripts were filtered according to the parameters of log2FC ≤ −0.3 and adjusted *P* value ≤ 0.05.

Functional analysis was performed in Cytoscape v3.6.1. Differentially expressed genes were grouped according to their gene ontology term using ClueGO v2.5.3 [20].

### Reverse transcription and quantitative PCR

2×10^5^ TCam-2 cells were transfected with 1.5 µg of constructs encoding the wild-type, mutated NANOS1 or empty pCMV6-entry vector in three biological replicates. Transcription was stopped by Actinomycin D (5 µg/ml) 24 h after transfection for 4 h. Total RNA isolation, template preparation and qPCR reactions conditions are described previously in [21]. Sequences of the primers are listed in Supplementary **Table S1**. *ARNT*, *GAPDH*, *RPL13* and *UBC* genes were used for normalization.

### Constructs

For the wild-type and mutant *NANOS1* construct overexpression, we used procedures previously described [22].

### Accession numbers

*NANOS1* NM_199461.4, *GADD45A* NM_001924.4, *GADD45B* NM_015675.4, *GADD45G* NM_006705.4, *RHOB* NM_004040.4, *BCL10* NM_003921.5, *STK17A* NM_004760.3, *TP53BP2* NM_001031685.3, *RIPK1* NM_003804.6, *SIAH1* NM_003031.4, *JUN* NM_002228.4.

### Statistical analysis

One-way unpaired t-test was used to estimate statistical significance. The *P* value < 0.05 (*) was considered statistically significant.

## Results

### Overexpression of NANOS1 containing the p.[(Pro34Thr);(Ser83del)] mutation results in reduced confluency and decreased cell proliferation

To understand the functional significance of the NANOS1 double mutation (**Fig. 1A**), especially its relationship to the absence of germ cells in seminiferous tubules in infertile male patients, we studied its effects on the phenotype of the TCam-2 cell line, which originates from human seminoma and represents male germ cells at an early stage of development [23]. While overexpression of the wild-type NANOS1 caused a significant increase (of nearly 50%) in cell confluency compared to the empty vector, overexpression of the mutated NANOS1 caused a significant decrease (of nearly 45%) in cell confluency compared to the empty vector (**Fig. 1B**). To check whether this negative effect on cell confluency reflected the influence of the NANOS1 mutation on cell proliferation, we overexpressed the wild-type and mutated NANOS1 in TCam-2 cells and performed MTS assay. We found that while the wild-type NANOS1 stimulated cell proliferation, the mutated NANOS1 caused a significant decrease in proliferation compared to the empty vector (**Fig. 1C**). We show that the NANOS1 mutation caused a functional switch of NANOS1 from pro-proliferative to anti-proliferative activity. The expression of both the NANOS1 constructs after transfection is shown in **Fig. 1D**.

### Overexpression of NANOS1 containing the p.[(Pro34Thr);(Ser83del)] mutation results in increased apoptosis

To determine whether the decreased cell proliferation upon overexpression of the NANOS1 mutant (**Fig. 1C**) was associated with increased apoptosis, we performed flow cytometry analysis of Annexin V-stained TCam-2 cells transfected with the wild-type or mutated NANOS1, as compared to empty vector. We found that while the wild-type NANOS1 reduced the spontaneous apoptosis of TCam-2 cells, the mutated NANOS1 induced it (**Fig. 2A**). Thus, we show that the NANOS1 mutation provoked a functional switch of NANOS1 from anti-apoptotic to pro-apoptotic. A quantification of the flow cytometric data is presented on the graph **Fig. 2B**, while the expression of the wild-type and mutated NANOS1 proteins in transfected cells are shown in **Fig. 2C**.

**Figure 2.**
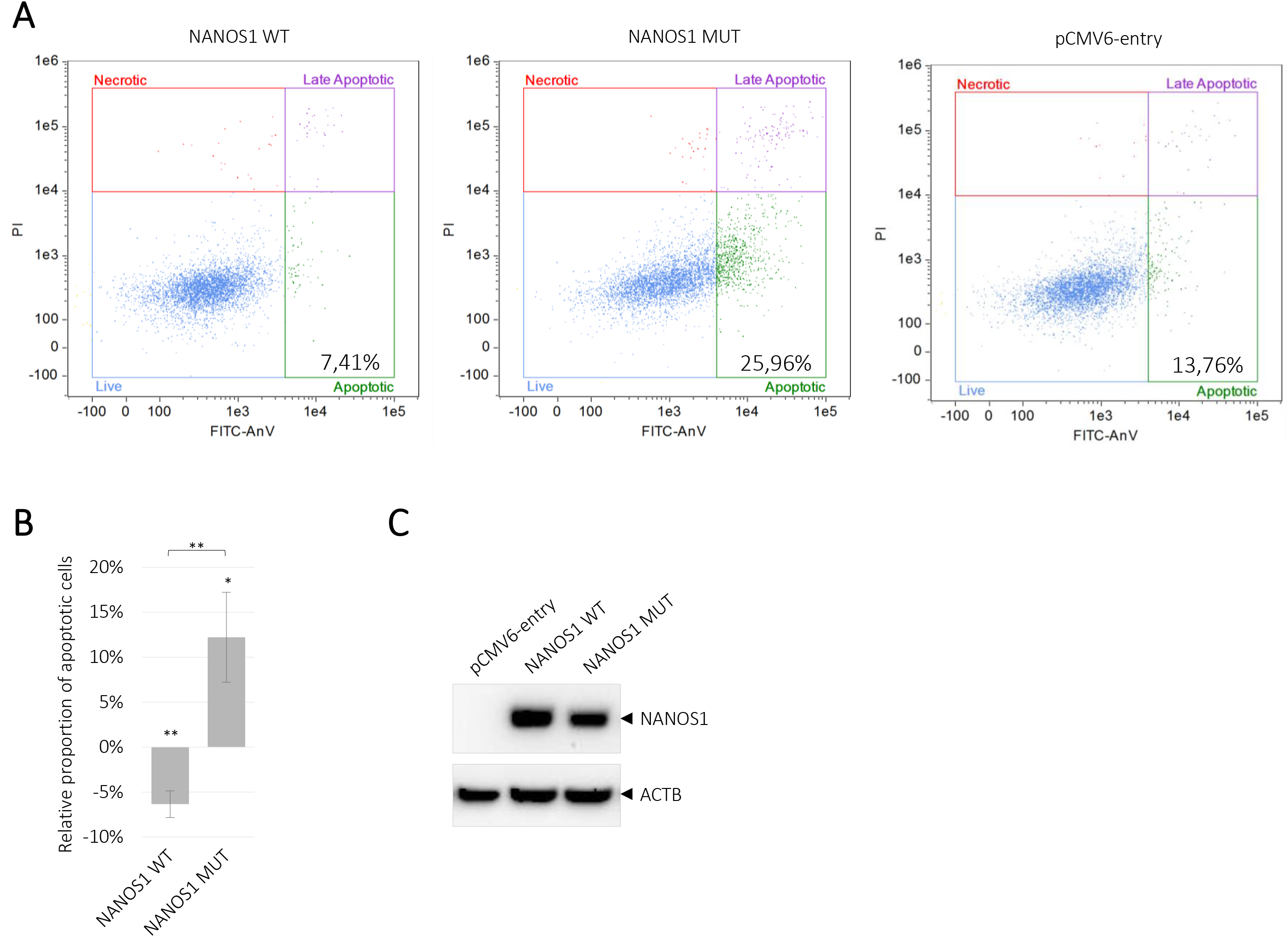
Analysis of the influence of the wild-type and mutated NANOS1 p.[(Pro34Thr);(Ser83del)] on apoptosis of TCam-2 cells. Apoptosis was measured by using Annexin V staining and flow cytometry. **A** – The representation of dot plots generated by flow cytometric analysis of apoptosis in TCam-2 cells upon transfection of the wild-type (WT), mutated (MUT) NANOS1 or empty pCMV6-entry plasmid. The percentages of apoptotic cells are shown in each plot. **B** – Quantitation of the data from the dot plots that represents the proportion of apoptotic cells in TCam-2 cells transfected with the wild-type or mutated NANOS1, relative to cells transfected with the empty vector (baseline). **C** – Western blot analysis after transfection of TCam-2 cells with pCMV6-entry vector, the wild-type or mutated NANOS1. As loading control was used ACTB (actin beta) protein, in each sample shown in lower panel. These experiment was performed in three biological replicates. The *P* value < 0.05 was considered statistically significant and was marked by one star (*) and the *P* value < 0.005 was marked by two stars (**).

### NANOS1-mediated repression of pro-apoptotic genes is disrupted by p.[(Pro34Thr);(Ser83del)] mutation

In order to identify apoptotic factors dysregulated by the p.[(Pro34Thr);(Ser83del)] NANOS1 mutation, first we performed a search for apoptosis-related transcripts regulated by wild-type NANOS1. Therefore, we carried out RNA-Seq analysis upon NANOS1 overexpression in TCam-2 cells. We identified 10 mRNAs encoding pro-apoptotic factors (*GADD45A*, *GADD45B*, *GADD45G*, *RHOB*, *BCL10*, *STK17A*, *TP53BP2*, *RIPK1*, *SIAH1*, *JUN*) which were downregulated upon wild-type NANOS1 overexpression (**Table S2**). Quantitative RT-PCR (RT-qPCR) confirmed downregulation of 7 among them (*GADD45A*, *GADD45B*, *GADD45G*, *RHOB, BCL10*, *STK17A*, *TP53BP2*) (**Fig. 3**). Finally, we investigated whether p.[(Pro34Thr);(Ser83del)] mutation influences that wild-type NANOS1-mediated repression effect by performing RT-qPCR upon overexpression of the mutated NANOS1. We found that the repression of 4 mRNAs (*GADD45A*, *GADD45B*, *GADD45G*, *RHOB*) was abrogated by the p.[(Pro34Thr);(Ser83del)] NANOS1 mutation (**Fig. 3**).

**Figure 3.**
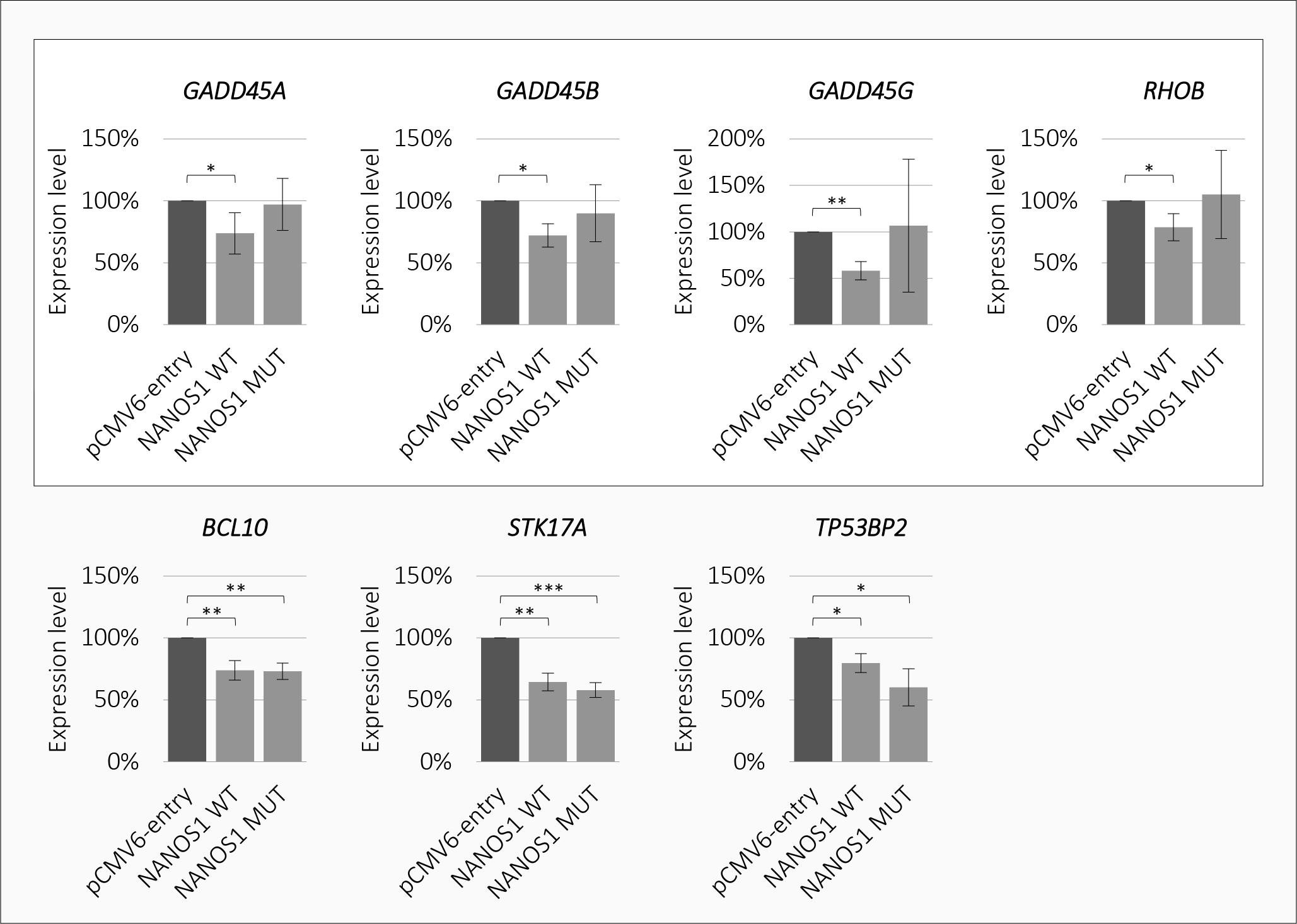
Quantitative RT-PCR analysis of selected pro-apoptotic genes in TCam-2 cells overexpressed the empty pCMV6-entry vector, the wild-type (WT) or mutated (MUT) NANOS1 p.[(Pro34Thr);(Ser83del)]. Repression of 7 mRNAs by the wild type NANOS1 is shown in the grey box, while disruption of repression of 4 mRNAs by the mutated NANOS1 is shown in the white box. The *P* value < 0.05 was considered statistically significant and was marked by one star (*), the *P* value <0.005 was marked by two stars (**) and the *P* value <0.0005 by three stars (***).

### The NANOS1 p.[(Pro34Thr);(Ser83del)] mutation does not affect NANOS1-mediated inhibition of cell cycle

Subsequently, we wanted to determine whether the differences in cell proliferation observed upon overexpression of the wild type or the mutated NANOS1 (**Fig. 1C**) were due to modulation of the cell cycle. To the best of our knowledge, quantification of cell distribution in cell cycle phases in response to Nanos has never been reported yet in any model. Therefore, we performed flow cytometric analysis of cell cycle distribution in unsynchronized TCam-2 cells overexpressing the wild-type or the mutated NANOS1, in comparison with the empty vector (**Fig. S1A**). The separation of cell populations in each cell cycle phase was performed using the ModFit program (**Fig. S1B**). Upon overexpression of the wild-type or the mutated NANOS1 we observed an increase in the proportion of cells in the G0/G1 phase, with a concordant decrease in the proportion of cells in the S phase, in comparison to the empty vector (**Fig. S1A**), indicating that NANOS1 inhibits TCam-2 cell cycle. However, no significant difference in cell cycle distribution was observed between cells expressing the wild-type and the mutated NANOS1 (**Fig. S1A**), This result indicates that this mutation does not alter the inhibitory effect of NANOS1 on cell cycle.

## Discussion

Previously, we have described a *NANOS1* gene p.[(Pro34Thr);(Ser83del)] mutation in infertile patients characterized by absence of germ cells in seminiferous tubules [14]. Here, we investigated the influence of this mutation on cellular processes which could account for that phenotype.

We showed for the first time that NANOS1 inhibits apoptosis and that the NANOS1 p.[(Pro34Thr);(Ser83del)] mutation caused increased apoptosis of TCam-2 cells, the human male germ line in culture. Although Nanos proteins have been reported as negative regulators of apoptotic process [4 5], molecular background of their anti-apoptotic properties is not known in mammals. Therefore, we hypothesised that NANOS1 negatively regulates apoptosis by repressing mRNAs encoding pro-apoptotic factors and that the p.[(Pro34Thr);(Ser83del)] mutation disrupts that regulation. Indeed, we identified 7 mRNAs encoding well-established pro-apoptotic factors which were repressed by NANOS1: *GADD45A*, *GADD45B*, *GADD45G*, *RHOB, BCL10*, *STK17A* and *TP53BP2*. Downregulation of these mRNAs by NANOS1 has never before been reported and may account for anti-apoptotic property of this protein in the human male germ cells. Importantly, we observed that *GADD45A*, *GADD45B*, *GADD45G* and *RHOB* mRNAs were not downregulated by the mutated NANOS1 p.[(Pro34Thr);(Ser83del)]. Interestingly, each of those four mRNAs encodes a protein activating DNA damage-induced apoptosis. Moreover, each of the three GADD45 (growth arrest and DNA damage inducible) protein family members induces apoptosis by binding and activating MAP3K4 kinase, which in turn positively regulates p38/JNK cascade [24]. RHOB (ras homolog family member B), on the other hand, activates apoptosis via BCL2L11 BIM activation [25]. Given our obtained results, we propose that lack of repression of above mentioned four pro-apoptotic mRNAs upon expression of p.[(Pro34Thr);(Ser83del)] NANOS1 mutation accounts for increased apoptosis of TCam-2 cells.

It is important to note that TCam-2, a human male germ cell line, originates from seminoma, which is a type of testis germ cell tumour [23]. In this context, it is interesting that almost all NANOS1-repressed pro-apoptotic factors identified in this study (GADD45A, GADD45B, GADD45G, RHOB and TP53BP2) are well established as tumour suppressors [26–34]. This is in line with NANOS1 itself being found to promote carcinogenesis [35]. Therefore, it is likely that in addition to playing a role in germ cell development, NANOS1 downregulates the expression of tumour suppressing factors, as has already been suggested for other RNA-binding proteins [36]. This finding represents an interesting issue for future studies aimed at investigating the role of NANOS1 in cancerogenesis.

Additionally, we showed that the wild-type NANOS1 caused an increase in the proportion of cells in the G0/G1 phase, with a concordant decrease of cells in the S phase. This is the first report, to the best of our knowledge, demonstrating cell cycle inhibition by quantification of cells in various phases of the cell cycle in response to Nanos protein. It is not clear why the mutation does not change the cell cycle effect caused by the wild-type protein, as it does in regard to the apoptosis effect. A potential changing effect of this mutation could be compensated by some other unknown factor(s) influencing the cell cycle in TCam-2 cells. However, addressing this question would require further investigation.

Altogether, the functional switch of NANOS1 from pro-proliferative towards anti-proliferative and from anti-apoptotic to pro-apoptotic, may represent the mechanism underlying the infertility in patients carrying the *NANOS1* p.[(Pro34Thr);(Ser83del)] mutation. In patients, this mutation could affect the regulation of specific mRNA targets such as *GADD45A*, *GADD45B*, *GADD45G* and *RHOB* at the post-transcriptional level, resulting in dysregulated expression of encoded factors necessary for germ cell maintenance, thereby causing the absence of germ cells in the seminiferous tubules, thus leading to infertility.

## Acknowledgements

We thank Dr. Sohei Kitazawa for providing us with the TCam-2 cell line and Dr. Riko Kitazawa for the maintenance of this cell line for us. This study was supported by grants from the National Science Centre Poland no 2014/15/B/NZ1/03384 to KKZ, no 2011/01/B/NZ2/04833 to BGM and ETIUDA scholarship no 2014/12/T/NZ1/00497 to MPS.

## References

1. Struhl G. Differing strategies for organizing anterior and posterior body pattern in Drosophila embryos. Nature 1989;338(6218):741–4 doi: 10.1038/338741a0[published Online First: Epub Date]∣.

2. Bhat KM The posterior determinant gene nanos is required for the maintenance of the adult germline stem cells during Drosophila oogenesis. Genetics 1999;151(4):1479–92

3. Asaoka-Taguchi M, Yamada M, Nakamura A, et al. Maternal Pumilio acts together with Nanos in germline development in Drosophila embryos. Nat Cell Biol 1999;1(7):431–7 doi: 10.1038/15666[published Online First: Epub Date]∣.

4. Sato K, Hayashi Y, Ninomiya Y, et al. Maternal Nanos represses hid/skl-dependent apoptosis to maintain the germ line in Drosophila embryos. Proc Natl Acad Sci U S A 2007;104(18):7455–60 doi: 10.1073/pnas.0610052104[published Online First: Epub Date]∣.

5. Suzuki A, Tsuda M, Saga Y. Functional redundancy among Nanos proteins and a distinct role of Nanos2 during male germ cell development. Development 2007;134(1):77–83 doi: 10.1242/dev.02697[published Online First: Epub Date]∣.

6. Jaruzelska J, Kotecki M, Kusz K, et al. Conservation of a Pumilio-Nanos complex from Drosophila germ plasm to human germ cells. Dev Genes Evol 2003;213(3):120–6 doi: 10.1007/s00427-003-0303-2[published Online First: Epub Date]∣.

7. Lolicato F, Marino R, Paronetto MP, et al. Potential role of Nanos3 in maintaining the undifferentiated spermatogonia population. Dev Biol 2008;313(2):725–38 doi: 10.1016/j.ydbio.2007.11.011[published Online First: Epub Date]∣.

8. Suzuki A, Saga Y. Nanos2 suppresses meiosis and promotes male germ cell differentiation. Genes Dev 2008;22(4):430–5 doi: 10.1101/gad.1612708[published Online First: Epub Date]∣.

9. Tsuda M, Sasaoka Y, Kiso M, et al. Conserved role of nanos proteins in germ cell development. Science 2003;301(5637):1239–41 doi: 10.1126/science.1085222[published Online First: Epub Date]∣.

10. Haraguchi S, Tsuda M, Kitajima S, et al. nanos1: a mouse nanos gene expressed in the central nervous system is dispensable for normal development. Mech Dev 2003;120(6):721–31

11. Lek M, Karczewski KJ, Minikel EV, et al. Analysis of protein-coding genetic variation in 60,706 humans. Nature 2016;536(7616):285–91 doi: 10.1038/nature19057[published Online First: Epub Date]∣.

12. Kusz K, Tomczyk L, Spik A, et al. NANOS3 gene mutations in men with isolated sterility phenotype. Mol Reprod Dev 2009;76(9):804 doi: 10.1002/mrd.21070[published Online First: Epub Date]∣.

13. Kusz KM, Tomczyk L, Sajek M, et al. The highly conserved NANOS2 protein: testis-specific expression and significance for the human male reproduction. Mol Hum Reprod 2009;15(3):165–71 doi: 10.1093/molehr/gap003[published Online First: Epub Date]∣.

14. Kusz-Zamelczyk K, Sajek M, Spik A, et al. Mutations of NANOS1, a human homologue of the Drosophila morphogen, are associated with a lack of germ cells in testes or severe oligo-astheno-teratozoospermia. J Med Genet 2013;50(3):187–93 doi: 10.1136/jmedgenet-2012-101230[published Online First: Epub Date]∣.

15. Bhandari D, Raisch T, Weichenrieder O, et al. Structural basis for the Nanos-mediated recruitment of the CCR4-NOT complex and translational repression. Genes Dev 2014;28(8):888–901 doi: 10.1101/gad.237289.113[published Online First: Epub Date]∣.

16. Bolger AM, Lohse M, Usadel B. Trimmomatic: a flexible trimmer for Illumina sequence data. Bioinformatics 2014;30(15):2114–20 doi: 10.1093/bioinformatics/btu170[published Online First: Epub Date]∣.

17. Langmead B, Salzberg SL. Fast gapped-read alignment with Bowtie 2. Nat Methods 2012;9(4):357–9 doi: 10.1038/nmeth.1923[published Online First: Epub Date]∣.

18. Liao Y, Smyth GK, Shi W. featureCounts: an efficient general purpose program for assigning sequence reads to genomic features. Bioinformatics 2014;30(7):923–30 doi: 10.1093/bioinformatics/btt656[published Online First: Epub Date]∣.

19. Love MI, Huber W, Anders S. Moderated estimation of fold change and dispersion for RNA-seq data with DESeq 2. Genome Biol 2014;15(12):550 doi: 10.1186/s13059-014-0550-8[published Online First: Epub Date]∣.

20. Bindea G, Mlecnik B, Hackl H, et al. ClueGO: a Cytoscape plug-in to decipher functionally grouped gene ontology and pathway annotation networks. Bioinformatics 2009;25(8):1091–3 doi: 10.1093/bioinformatics/btp101[published Online First: Epub Date]∣.

21. Janecki DM, Sajek M, Smialek MJ, et al. SPIN1 is a proto-oncogene and SPIN3 is a tumor suppressor in human seminoma. Oncotarget 2018;9(65):32466–77 doi: 10.18632/oncotarget.25977[published Online First: Epub Date]∣.

22. Sajek M, Janecki DM, Smialek MJ, et al. PUM1 and PUM2 exhibit different modes of regulation for SIAH1 that involve cooperativity with NANOS paralogues. Cell Mol Life Sci 2019;76(1):147–61 doi: 10.1007/s00018-018-2926-5[published Online First: Epub Date]∣.

23. de Jong J, Stoop H, Gillis AJ, et al. Further characterization of the first seminoma cell line TCam-2. Genes Chromosomes Cancer 2008;47(3):185–96 doi: 10.1002/gcc.20520[published Online First: Epub Date]∣.

24. Takekawa M, Saito H. A family of stress-inducible GADD45-like proteins mediate activation of the stress-responsive MTK1/MEKK4 MAPKKK. Cell 1998;95(4):521–30 doi: 10.1016/s0092-8674(00)81619-0[published Online First: Epub Date]∣.

25. Marlow LA, Bok I, Smallridge RC, et al. RhoB upregulation leads to either apoptosis or cytostasis through differential target selection. Endocr Relat Cancer 2015;22(5):777–92 doi: 10.1530/ERC-14-0302[published Online First: Epub Date]∣.

26. Adnane J, Muro-Cacho C, Mathews L, et al. Suppression of rho B expression in invasive carcinoma from head and neck cancer patients. Clin Cancer Res 2002;8(7):2225–32

27. Liu B, Yang L, Li XJ, et al. Expression and significance of ASPP2 in squamous carcinoma of esophagus. Kaohsiung J Med Sci 2018;34(6):321–29 doi: 10.1016/j.kjms.2017.12.011[published Online First: Epub Date]∣.

28. Mazieres J, Antonia T, Daste G, et al. Loss of RhoB expression in human lung cancer progression. Clin Cancer Res 2004;10(8):2742–50

29. Mukherjee K, Sha X, Magimaidas A, et al. Gadd45a deficiency accelerates BCR-ABL driven chronic myelogenous leukemia. Oncotarget 2017;8(7):10809–21 doi: 10.18632/oncotarget.14580[published Online First: Epub Date]∣.

30. Prendergast GC. Farnesyltransferase inhibitors define a role for RhoB in controlling neoplastic pathophysiology. Histol Histopathol 2001;16(1):269–75 doi: 10.14670/HH-16.269[published Online First: Epub Date]∣.

31. Sha X, Hoffman B, Liebermann DA. Loss of Gadd45b accelerates BCR-ABL-driven CML. Oncotarget 2018;9(70):33360–67 doi: 10.18632/oncotarget.26076[published Online First: Epub Date]∣.

32. Song B, Bian Q, Zhang YJ, et al. Downregulation of ASPP2 in pancreatic cancer cells contributes to increased resistance to gemcitabine through autophagy activation. Mol Cancer 2015;14:177 doi: 10.1186/s12943-015-0447-5[published Online First: Epub Date]∣.

33. Tian L, Deng Z, Xu L, et al. Downregulation of ASPP2 promotes gallbladder cancer metastasis and macrophage recruitment via aPKC-iota/GLI1 pathway. Cell Death Dis 2018;9(11):1115 doi: 10.1038/s41419-018-1145-1[published Online First: Epub Date]∣.

34. Wu T, Song H, Xie D, et al. Silencing of ASPP2 promotes the proliferation, migration and invasion of triple-negative breast cancer cells via the PI3K/AKT pathway. Int J Oncol 2018;52(6):2001-10 doi: 10.3892/ijo.2018.4331[published Online First: Epub Date]∣.

35. Bonnomet A, Polette M, Strumane K, et al. The E-cadherin-repressed hNanos1 gene induces tumor cell invasion by upregulating MT1-MMP expression. Oncogene 2008;27(26):3692–9 doi: 10.1038/sj.onc.1211035[published Online First: Epub Date]∣.

36. Blackinton JG, Keene JD. Post-transcriptional RNA regulons affecting cell cycle and proliferation. Semin Cell Dev Biol 2014;34:44–54 doi: 10.1016/j.semcdb.2014.05.014[published Online First: Epub Date]∣.

