## Supplemental Fig.S1 for "A male infertility mutation reverts NANOS1 activity from anti-apoptotic to pro-apoptotic by disrupting repression of *GADD45A*, *GADD45B*, *GADD45G* and *RHOB* genes"

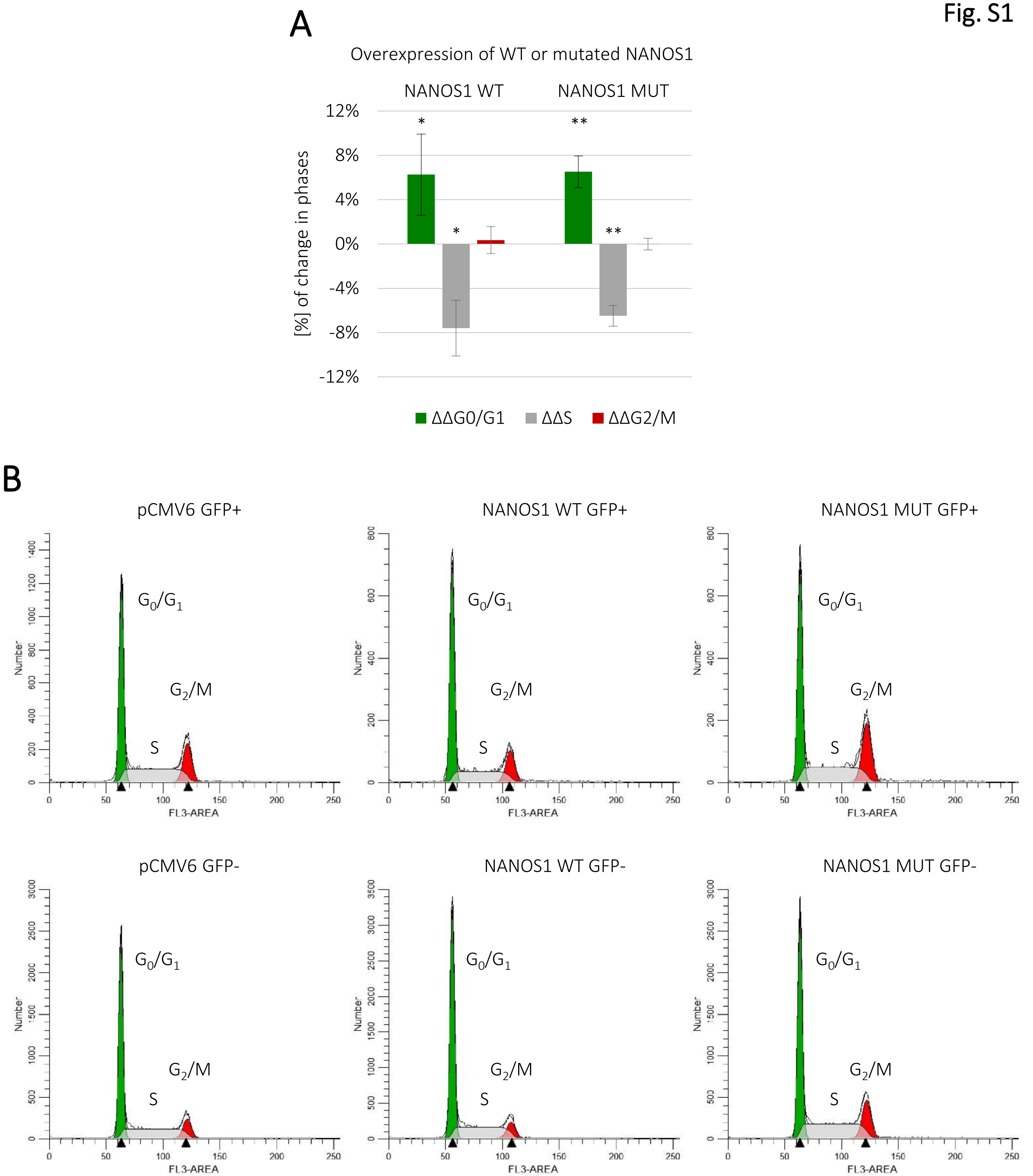


**Figure S1.** Cell cycle analysis of propidium iodide stained TCam-2 cells by flow cytometry. **A**. The graph represents distribution at G0/G1, S and G2/M cell cycle phases of unsynchronized TCam-2 cells overexpressing the wild-type (WT) or mutated (MUT) NANOS1 p.[(Pro34Thr);(Ser83del)] compared to the empty pCMV6-entry control vector (baseline). Flow cytometry measurement of the distribution was performed 48 h after transfection with construct encoding the wild-type, the mutated NANOS1 or the empty pCMV6-entry co-transfected with GFP-F. Percentages of cells in each cell cycle phase were calculated using ModFit LT software. This experiment was repeated in four independent experiments. The *P* value < 0.05 was considered statistically significant and was marked by one star (*) and the *P* value < 0.005 was marked by two stars (**).**B.** Histogram representation of the TCam-2 cell distribution at G0/G1, S and G2/M phases of the cell cycle in unsynchronized culture upon transfection with construct encoding the wild-type (WT) or mutated (MUT) NANOS1 or the empty pCMV6-entry vector co-transfected with GFP-F. Histograms were generated by ModFit LT software and show one representative example from four independent experiments.
