## Supplemental Tables for "A male infertility mutation reverts NANOS1 activity from anti-apoptotic to pro-apoptotic by disrupting repression of *GADD45A*, *GADD45B*, *GADD45G* and *RHOB* genes"

**Supplementary Table S1.** List of primers used for RT-qPCR. For primers, F indicates forward and R – revers.

| Gene | Primer | Sequence 5′ → 3′ | Primer length [nt] | Amplicon length [nt] | Annealing temperature [°C] |
|---|---|---|---|---|---|
| BCL10 | F | AATACCATCTTCTCTTCA | 18 | 91 | 56 °C |
| | R | TTCCTTCTTCTTCTAACT | 18 | | |
| GADD45A | F | GTGACGAATCCACATTCATCTC | 22 | 90 | 56 °C |
| | R | CCATTGATCCATGTAGCGACTT | 22 | | |
| GADD45B | F | AAGTTGATGAATGTGGAC | 18 | 135 | 56 °C |
| | R | GATGTTGATGTCGTTGTC | 18 | | |
| GADD45G | F | TCAGCCAAAGTCTTGAAC | 18 | 107 | 56 °C |
| | R | ATCAGCGTAAAATGGATCT | 19 | | |
| RHOB | F | TGCTGATCGTGTTCAGTAAG | 20 | 91 | 56 °C |
| | R | TTGCCGTCCACCTCAATG | 18 | | |
| STK17A | F | TGAACTAGCACAAGACAATCCT | 22 | 91 | 56 °C |
| | R | AGCAGCATATTCCAGAACTAAGA | 23 | | |
| TP53BP2 | F | ACCAGAGCAGTGAAGATA | 18 | 135 | 56 °C |
| | R | CTGAAGGTGGCTGATTAG | 18 | | |
| ARNT | F | CCACAG-GAACTCTTAGGAA | 19 | 117 | 56 °C |
| | R | CATGACAGACAGCACTTG | 18 | | |
| GAPDH | F | CGGAGTCAACGGATTTGGTCGTAT | 24 | 307 | 56 °C |
| | R | AGCCTTCTCCATGGTGGTGAAGAC | 24 | | |
| UBC | F | ATTTGGTGCGCGGTTCTTG | 19 | 114 | 56 °C |
| | R | TGCCTTGACATTCTCGATGGT | 21 | | |
| RPL13 | F | CCTGGAGGAGAAGGAGGGAAAGAGA | 25 | 102 | 56 °C |
| | R | TTGAGGACCTCTGTGTATTTGTCAA | 25 | | |

**Supplementary Table S2.** List of pro-apoptotic genes selected from RNA-Seq analysis as downregulated by NANOS1. Expression of these genes were reduced in TCam-2 cells transfected with the wild type NANOS1 construct in comparison to cells transfected with the empty vector.

| HGNC | Base Mean | log2 Fold Change | Adjusted *P* value | *P* value | Standard Error |
|---|---|---|---|---|---|
| GADD45A | 1051,99 | -0,55 | 8,22E-04 | 3,93E-05 | 0,13 |
| GADD45B | 311,44 | -0,65 | 6,91E-04 | 3,08E-05 | 0,16 |
| GADD45G | 334,60 | -0,64 | 4,47E-03 | 3,97E-04 | 0,18 |
| RHOB | 516,17 | -0,42 | 3,00E-02 | 5,78E-03 | 0,15 |
| BCL10 | 32,64 | -0,64 | 3,32E-02 | 6,68E-03 | 0,24 |
| STK17A | 392,07 | -0,62 | 7,51E-04 | 3,48E-05 | 0,15 |
| TP53BP2 | 894,20 | -0,51 | 4,09E-04 | 1,57E-05 | 0,12 |
| RIPK1 | 294,70 | -0,41 | 3,69E-02 | 7,70E-03 | 0,15 |
| SIAH1 | 26,98 | -0,78 | 9,31E-03 | 1,11E-03 | 0,24 |
| JUN | 234,83 | -0,62 | 1,63E-03 | 1,00E-04 | 0,16 |
